# Diversity begets diversity in human cultures and mammal species

**DOI:** 10.1101/2020.04.28.066969

**Authors:** Marcus J. Hamilton, Robert S. Walker, Chris Kempes

## Abstract

A key feature of the distribution of life on Earth is the positive correlation between environmental productivity and biodiversity. This correlation also characterizes the distribution of human cultural diversity, which is highest near the equator and decreases exponentially toward the poles. Moreover, it is now understood that the tropics house more biodiversity than would be expected from energy availability alone suggesting “diversity begets diversity”. Here we show the same is also true for human cultural diversity. This convergence is particularly striking because while the dynamics of biological and cultural evolution may be similar in principle the mechanisms and time scales involved are very different. However, a common currency underlying all forms of diversity is ecological kinetics; the temperature-dependent fluxes of energy and biotic interactions that sustain life at all levels of biological and social organization. Using macroecological theory and the analysis of global databases we show both mammal diversity and cultural diversity scale superlinearly with environmental productivity at rates predicted by the ecological kinetics of environmental productivity. Diversity begets diversity in human cultures and mammal species because the kinetics of energy availability and biotic interactions result in superlinear scaling with environmental productivity.

## Introduction

Biodiversity and cultural diversity are distributed unequally across the planet’s surface [1–7], as characterized by one of the most pervasive features of the biogeographic distribution of life on Earth, the latitudinal gradient of biodiversity [8–10]. Across taxa, from mammals, birds, reptiles, and amphibians [11,12], to plants [13–15], marine bivalves [16], ants [17], liverworts [18], and diseases [19], species richness is highest at the equator and decays exponentially toward the poles. These biodiversity gradients extend deep into the geological past [8,9,20,21]. Human cultural, linguistic, and economic diversity show the same gradients [1,3,4,22,23]. While there is no unified mechanistic theory of biogeography to explain these biodiversity gradients, proposed hypotheses tend to focus on some combination of ecological (i.e., niche diversity), energetic (i.e., ecosystem kinetics), evolutionary (i.e., phylogenetic history), and/or historical mechanisms (i.e., dispersal) [11,13,24–28]. For cultural diversity, similar mechanisms have been proposed, including latitudinal gradients in the length of the growing season [22,23], environmental gradients [1,4,29], and the timing of initial human colonization of continents [2].

While latitudinal gradients of diversity are pervasive, latitude itself cannot be explanatory as it is simply a geographic binning procedure that captures the steady loss biodiversity with increasing distance from the equator. Latitude indexes large-scale variation at many scales including the intensity of solar radiation, the length of the growing season, the distribution of precipitation, and the statistics of environmental temperature, all of which interact to constrain the distribution of environmental productivity around the planet. Exactly how the productivity of local environments influences biodiversity has been central to ecological and evolutionary theory since Darwin and Wallace [30].

While the observation is commonly made that the global distributions of biodiversity and cultural diversity are remarkably similar, few studies have asked why [3,4,6]. At a fundamental level diversity must be correlated with the flux of energy through ecosystems as all organisms compete for environmental production to fuel metabolic processes of growth, maintenance and reproduction. Moreover, ecosystems differ in their capacity to support life due to environmental and climatic constraints on metabolic fluxes of free energy and so the availability of energy in local environments constrains total biomass, organismal abundance, and species diversity [25,31]. Thus, the biogeographic distribution of biodiversity covaries with geographic variation in energy availability. Despite technological innovations that enhance the ability of humans to compete for resources by reshaping their environments and redistributing materials, energy and information at multiple scales, ultimately the biogeographic distribution of human biomass is similarly constrained by geographic variation in energy availability [32–36].

Much classic theory in ecology proposed that latitudinal biodiversity gradients result from a direct link between environmental production and species diversity; more production supports more individuals that belong to more species, a.k.a. the “more-individuals-hypothesis” [24,37,38]. It is increasingly apparent, however, that species diversity is not simply a linear function of environmental productivity as the link between diversity and environmental productivity is more complex [25]. For example, while more productive environments house more species, tropical ecosystems house more diversity at all scales than would be expected by environmental production alone [14,25]. Similarly, studies of human cultures show that while diversity tends to be higher in more productive environments [1,3,4,6,22,23,39] there are many other environmental, climatic, geological, and cultural processes that impact diversity at various scales [39–44]. For example, while variation in global hunter-gatherer population density, space use, and mobility is well-predicted by environmental productivity [43,45,46], the spatial extent of ethnolinguistic groups across the socioeconomic spectrum is better predicted by the level of sociopolitical complexity [44]. Within continents, statistical models vary in their predictive power over space, presumably because local information becomes increasingly necessary to make predictions at finer scales of resolution; because the world is a scale-dependent complex system there can be no single predictor that explains diversity at all scales [see 37].

In this paper, we use these studies as motivation to examine the macroecological relationship between mammal species diversity and cultural diversity through the lens of the metabolic theory of ecology. Our goal in this paper is to explain why human cultural and mammalian diversity are so similarly distributed at the global scale and why both forms of diversity increase faster than would be predicted from environmental production alone.

### Individual metabolism

We start by considering the link between individual and ecosystem level metabolism. The basal metabolic rate of an individual organism, *B*_*i*_, can be expressed in terms of body mass, *M*, as

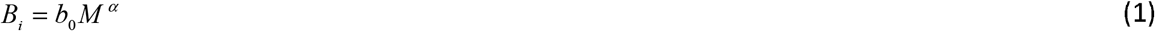

where *b*_0_ is a taxon-specific constant, independent of mass and temperature, and *α* ≈ 3/ 4, known as Kleiber’s Law [47]. This 3/4-power scaling of mass and metabolism arises from the fractal-like, space-filling nature of optimized internal networks that distribute the energy and resources that sustain organisms (such as the vascular system) [48–50]. The overall scale parameter, *b*_0_, is derived from the underlying biochemical kinetics of metabolism and, as such, depends on the temperature, *T*, at which an organism operates [50]. This is given by the exponential Arrhenius-Boltzmann factor, exp(−*E*_*B*_/*kT*), where *E*_*B*_ (~0.65 eV) is the average activation energy of the biochemical reactions contributing to metabolic processes of respiration, *k* is Boltzmann’s constant (8.62×10^−5^ eV K^−1^), and *T* is the absolute temperature at which the organism operates (°K) [51]. For endotherms, *T* is the internal body temperature, and for ectotherms, *T* is the ambient environmental temperature. Incorporating this temperature dependence into equation 1 yields the expression

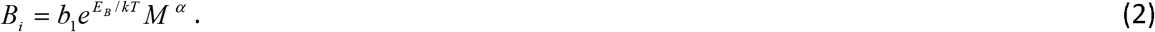

which is the central equation of metabolic theory of ecology [50–52].

### Ecosystem metabolism

As ecosystems are composed of individual organisms, it follows that the respiration of an entire ecosystem is the sum over all individuals, the majority of which are microbes and plants [53]. Temperature is one of the primary drivers of energy flux in ecosystems [54]. The largest ecological flux is gross primary production, *GPP*, the total mass of carbon entering an ecosystem through the conversion of solar radiation into biomass by photosynthesis over a given period of time [55,56]. At steady state, about half of *GPP* is respired by plants to support production, maintenance and ion uptake [57]. The remainder is used by plants to fuel growth, and this is terrestrial net primary production (*NPP*). *NPP* is thus the net carbon gain in an ecosystem over a given period of time measured in units of g C m^−2^ yr^−1^. Following mass-energy equivalence, *NPP* is thus the total amount of ecosystem energy available for work (i.e., Gibbs free energy). It then follows the flux of free energy in ecosystems is captured by the Arrhenius-Boltzmann temperature-dependence of *NPP*; 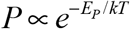, [53,58]. Here, if *E*_*P*_ is assumed to be the activation energy of photosynthesis, then holding all other ecological and evolutionary processes constant *E*_*P*_ ≈ 0.3 eV. However, all is not constant as the kinetics of *NPP* responds to other rate limiting constraints such as stoichiometry, climatic constraints, and water-availability [25,31,54,59]. Here we measure the temperature-dependence of *NPP* from data: introducing annual precipitation, *R* (mm yr^−1^), as a rate-limiting constraint on the kinetics of *NPP*, *P*, we have

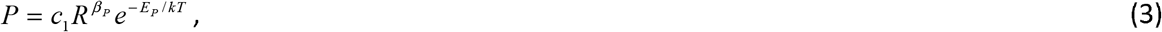

where *β*_*P*_ is an exponent capturing the effect of precipitation on *NPP* and *c*_1_ is a normalization constant independent of precipitation or temperature. We fit a multiple regression model of the form of equation 3 to our data and present the results in Figure 1. Results show global scale variation in *NPP* is well-predicted by temperature and precipitation (*R*^2^ = 92%), where the kinetics have a slope *E*_*Y*_ = 0.52 and precipitation is a sublinear constraint on *NPP*, *β*_*Y*_ = 0.72. Figure 1 shows a 3D surface plot of NPP as a function of temperature and precipitation.

**Figure 1.**
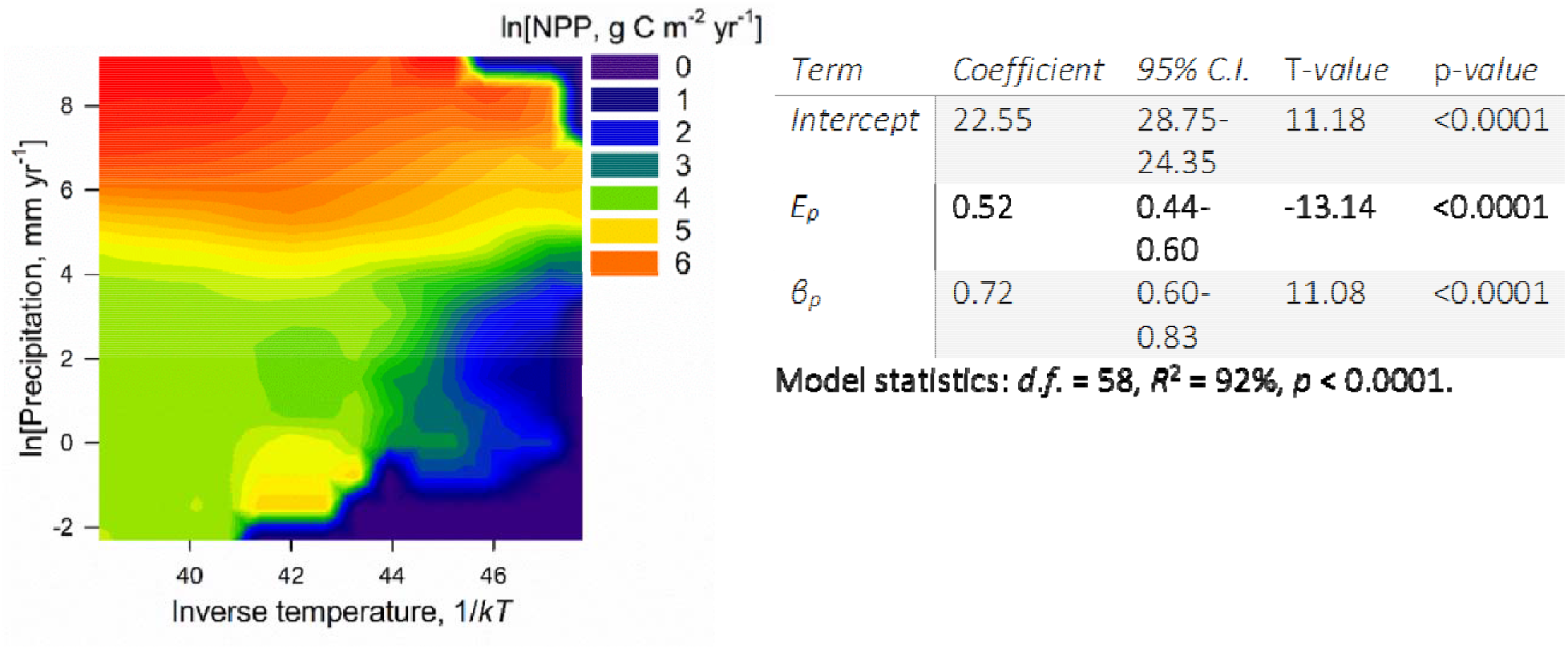
3D surface plot of net primary production as a function of temperature and precipitation in our data. To the right are the results of a multiple regression model of NPP as function of temperature and precipitation.

### Ecosystem complexity and biodiversity

While energy availability within ecosystems is ultimately determined by temperature-dependent fluxes of carbon, increasingly productive environments are also more structurally complex. That is to say, a polar desert is not simply an energy-poor tropical rainforest, and a tropical rainforest is not simply an energy rich grassland. Increasingly productive environments not only have greater energy flux per unit area but also house more species [25]. Evolutionary explanations suggest speciation and extinction dynamics vary consistently across environments affecting the standing stock of diversity, and data suggest the tropics are more taxonomically and genetically diverse [60,61]. Niche-based explanations suggest that in productive environments resources are apportioned in increasingly different ways, thus increasing the number of species that may co-exist in an ecosystem, and phylogenetically-related species may have a tendency to inhabit similar niches in similar environments [62–64]. Ecological speciation explanations suggest increasing interactions between species in more productive environments promote enhanced diversity through increased competition and constraints on dispersal. The metabolic theory of ecology suggests a more general kinetic explanation, not necessarily exclusive of the preceding theories; because the pace of life is driven by metabolic processes, all the ecological and evolutionary dynamics that constitute the structure of ecosystems are governed by biological kinetics where biotic interactions of all kinds occur faster when the environment is warmer [25,50]. Diversity increases both as a function of more energy and faster biotic interactions in more productive environments.

Because humans are also necessarily embedded within ecosystems, the abundance, density, and the intraspecific diversity of the human species are similarly subject to ecological kinetics.

Groups of human compete with each other for space and energy, as well as with other species, and cultural diversity is the result of intraspecific group competition and multiple other evolutionary processes playing out over time [65–67]. Here, we use ethnolinguistic diversity as our measure of cultural diversity. The term ethnolinguistic refers to a spatially and culturally discrete population of language speakers, which may or may not share a common language with other groups around the planet, such as English speakers in the British Isles, Australia, or the Caribbean, or Diné speakers in Alaska or the American Southwest. In both of these cases, each language is composed of multiple ethnolinguistic populations.

We model the kinetics of diversity by assuming 1) that the number of species increases with environmental productivity, and 2) as the number of species increases there are more interactions between those species. Therefore, we write the kinetics of diversity as a function of both the kinetics of energy flux and all other biotic interactions:

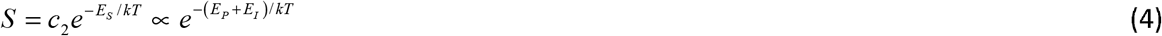

where *E*_*S*_ is the overall kinetics of diversity and *E*_*I*_ is the kinetics of ecological interactions. Here, the kinetics of diversity, *E*_*S*_, will be faster than the kinetics of ecosystem energy flux, *E*_*P*_, when the kinetics of biological interactions *E*_*I*_ > 0 . From equations 3 and 4 we can express species diversity, *S*, as a function of environmental productivity, *P*, as

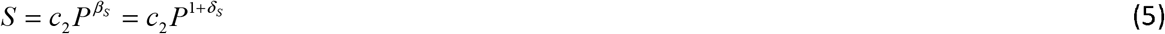

where *β*_*S*_ = *E*_*S*_ / *E*_*P*_ = (*E*_*P*_ + *E*_*I*_) / *E*_*P*_ = 1+ *δ*_*S*_, and so *δ*_*S*_ = *E*_*I*_ / *E*_*P*_. Therefore, diversity will increase superlinearly (*β*_*S*_ > 1) with environmental productivity whenever the kinetics of diversity are faster than the kinetics of environmental productivity (*E*_*S*_ > *E*_*P*_). When there are no interaction kinetics (*E*_*I*_ = 0), diversity will increase linearly with environmental productivity, and when interaction kinetics are negative (*E*_*I*_ < 0) then diversity will increase sublinearly. This latter case could result from competitive exclusion, for example.

To establish whether this simple model of ecological kinetics captures variation in biodiversity and human cultural diversity we use data on the global distribution of mammal species and ethnolinguistic populations to test four hypotheses using predictions derived from this model:

H_1_: **Diversity is a function of kinetics**. We first test the hypothesis that diversity increases exponentially with temperature, as predicted by ecological theory at a rate consistent with the kinetics of environmental metabolism.
H_2_: **Diversity kinetics is faster than productivity kinetics**. The kinetics of diversity, *E*_*S*_, should be faster than the kinetics of environmental productivity, *E*_*P*_, as species interaction rates, *E*_*I*_, are predicted to be both non-zero and increase with environmental productivity, *P* .
H_3_: **Diversity is more than environmental productivity**. Diversity, *S*, should increase with environmental productivity, *P*, at a superlinear rate given by *β*_*S*_ = *E*_*S*_ / *E*_*P*_ > 1.
H_4_: **Diversity kinetics are faster in more productive environments**. Holding NPP constant, diversity should increase with temperature, i.e., for a given range of NPP relatively warmer environments will be more diverse, and the slope of this response should increase in more productive environments.

## Results

### Hypotheses 1 and 2: Exponential latitudinal and temperature gradients

Figure 2 shows that the spatial distribution of mammal species (red points) and ethnolinguistic diversity (blue points) across the surface of the planet are similar. Inset along the Y-axis of Figure 2 are distributions of the number of ethnolinguistic groups and the number of mammal species by latitude. The two distributions show a high degree of overlap and a *t*-test shows they are statistically indistinguishable (*t*-test: *T*-value = −0.38, *d.f*. = 241, *P* = 0.701). Moreover, Figure 3A shows that after controlling for available landmass ethnolinguistic diversity and mammal diversity exhibit similar responses to absolute latitude. Exponential functions fit to the data shows ethnolinguistic diversity decreases with absolute latitude at a rate 0.05 ± 0.004 (95% C.I.) (OLS regression: *d.f*. = 73, *r*^2^ = 83%, *p* < 0.0001) and mammalian diversity as 0.05 ± 0.01 (OLS regression: *d.f*. = 68, *r*^2^ = 76%, *p* < 0.0001); both forms of diversity decrease at a constant rate of ~5% per degree latitude, and the correlation coefficient between the two data sets is 0.84 (Figure 3A inset).

**Figure 2.**
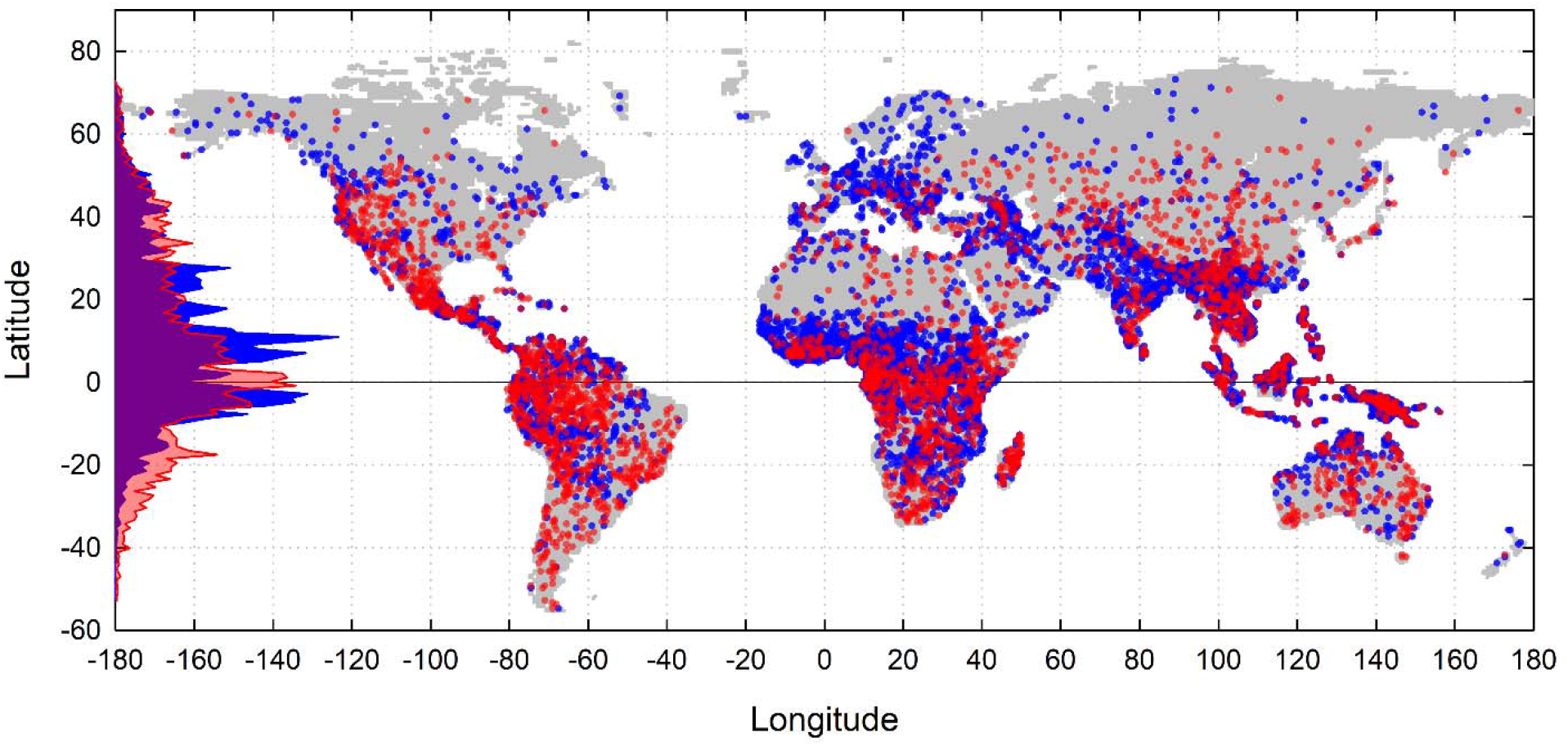
A map of the global distribution of human ethnolinguistic groups (blue) and mammal species (red). The underlying map is composed of the ~57,000 cells (0.5*0.5-degree latitude grids) comprising the global sampling procedure. Latitudinal frequency distributions of diversity are shown vertically along the Y-axis. Both forms of diversity are highest near the equator and decay toward the poles. However, both distributions show a frequency trough along the equator where ethnolinguistic diversity is highest approximately ±5 degrees from the equator, and mammal species diversity peaks just south of the equator.

**Figure 3.**
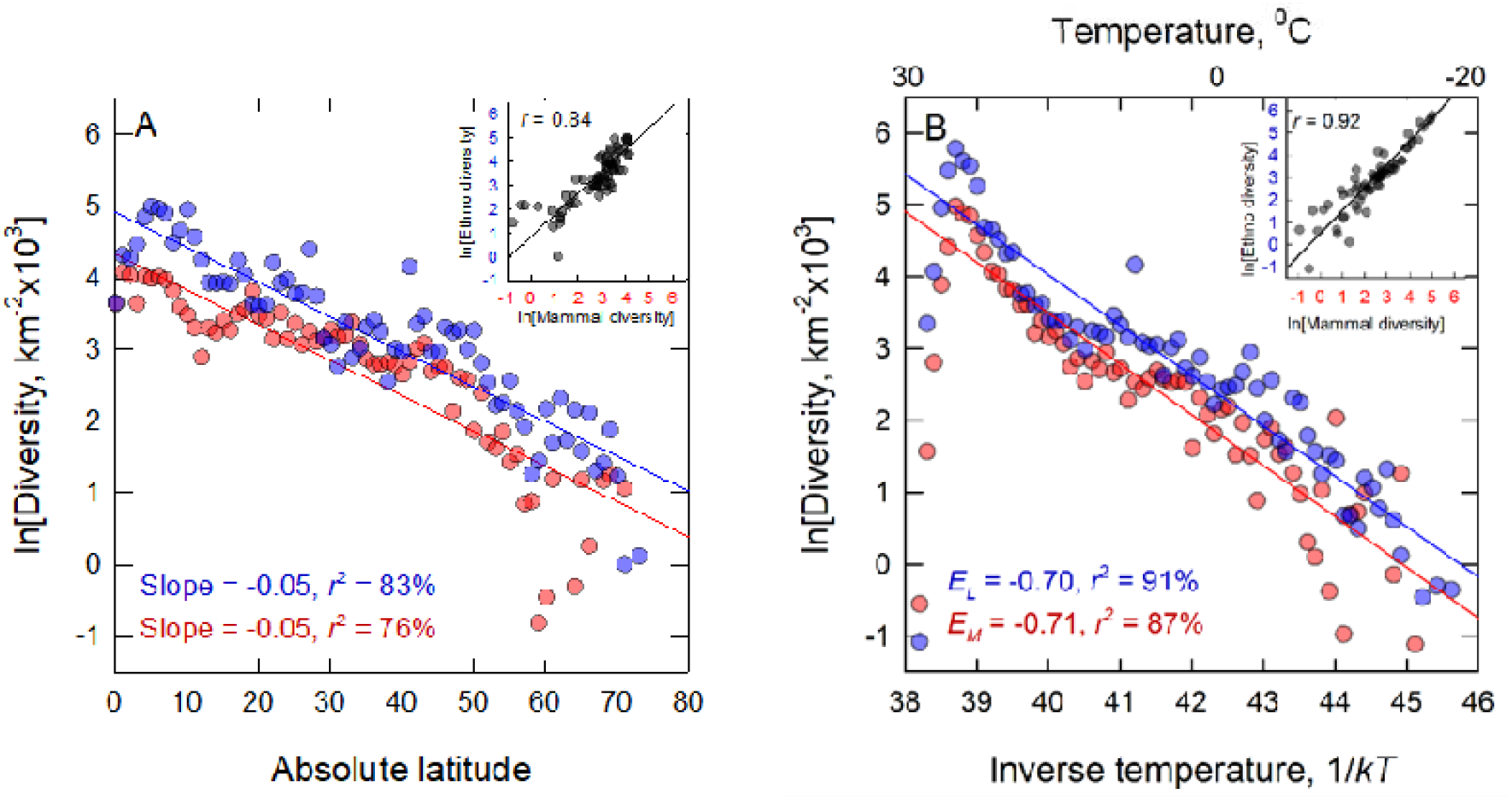
Latitudinal gradients (A) and temperature gradients (B) in ethnolinguistic diversity (blue points) and mammal species diversity (red points). In both cases both forms of diversity decrease exponentially with latitude and temperature at statistically indistinguishable rates. Inset figures are the correlations between the two forms of diversity.

In support of hypothesis 1, Figure 3B shows the exponential relationship between diversity and environmental temperature (in ^0^C along the upper *x*-axis and inverse temperature 1/*kT* along the lower *x*-axis) in both ethnolinguistic and mammalian diversity. A regression model of the form *y* = *y*_0_ exp(*E* / *kT*) is fit to both data sets. Ethnolinguistic diversity decreases with temperature at a rate *E*_*L*_ = −0.70 (0.65−0.76 95% C.L.) (OLS regression: *d.f.* = 65, *r*^2^=91%, p<0.0001). Similarly, mammal species diversity decreases with temperature at a rate *E*_*M*_ = −0.71 (0.64 − 0.78) (OLS regression: *d.f*. = 60, *r*^2^=87%, *p*<0.0001: for both data sets, statistical fits excluded the first three leftmost data points as they are outliers and have undue influence on the fit of the slope). The mammal and ethnolinguistic slopes are not significantly different and the two forms of diversity are highly correlated in space; *r* = 0.92 (inset figure 3B). The slopes of both the mammal and ethnolinguistic group fits are significantly steeper the slope of NPP shown in Figure 1 (*E*_*P*_ = 0.52), therefore providing support for hypothesis 2: The kinetics of diversity are steeper than the kinetics of productivity.

### Hypothesis 3: Diversity and net primary production superlinear scaling

Figure 4 shows tha both forms of diversity scale superlinearly with net primary production, thus providing support for hypothesis 3. Ethnolinguistic diversity increases superlinearly with net primary production, diversity as *β*_*L*_ = 1.40 (1.25 −1.55) (OLS regression: *d.f*. = 40, *r*^2^ = 91%, *p* < 0.0001) and mammalian diversity increases as *β*_*M*_ = 1.23 (1.10 −1.35) (OLS regression: *d.f*. = 36, *r*^2^ = 91%, *p* < 0.0001) (fits exclude the rightmost data point as it is an outlier and has a large influence of the fit of the slope). Both slopes are significantly greater than 1 and the data are highly correlated; *r* = 0.97 (inset figure). Moreover, the confidence intervals around the slopes both encompass their predicted values, thus providing support for hypothesis 4. For mammals, the predicted slope based on the ratio of the kinetic exponents is *λ*_*M*_ = *E*_*M*_ / *E*_*P*_ = 0.70 / 0.52 = 1.35 and the observed confidence interval is 1.10-1.35. For ethnolinguistic groups the predicted slope is *λ*_*L*_ = *E*_*L*_ / *E*_*P*_ = 0.71/ 0.52 = 1.37, and the observed confidence interval is 1.25-1.55.

**Figure 4.**
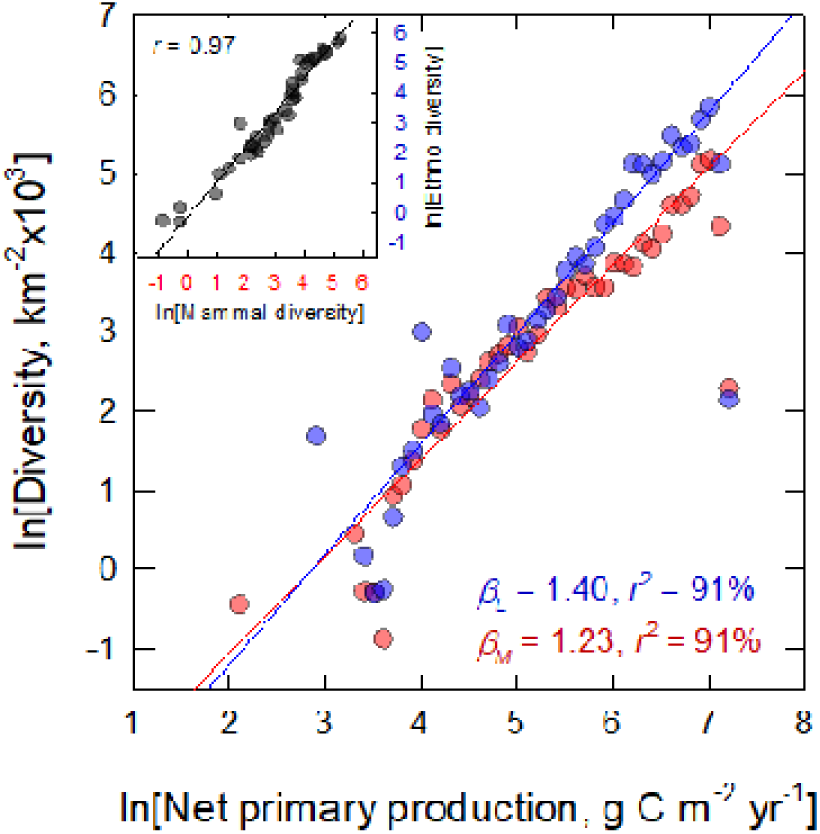
Ethnolinguistic and mammalian diversity per thousand square kms by (A) temperature and (B) net primary production. Inset figures are the resulting correlation plots of ethnolingistic and mammalian diversity. Solid lines are OLS regression fits, blue = ethnolinguistic and red = mammal.

### Hypothesis 4: Diversity and kinetics within NPP bins

Figures 5A and 5B show that both forms of diversity increase exponentially with temperature at faster rates in more productive environments. Here the diversity data are aggregated into bins of width 1lnNPP and exponential functions are fit to the data within each bin. Details of the regression models and the statistical results are given in the Supplementary Information. The inset figures are plots of the slope of the fit within each NPP bin by the intercept. Both inset panels show that the gradient of the slope increases by 0.02eV with each additional species or language. Therefore, these figures demonstrate that for both ethnolinguistic groups and mammal species diversity indeed begets diversity as the interaction rates in more productive environments increase with increasing diversity, thus providing support for hypothesis 4.

**Figure 5.**
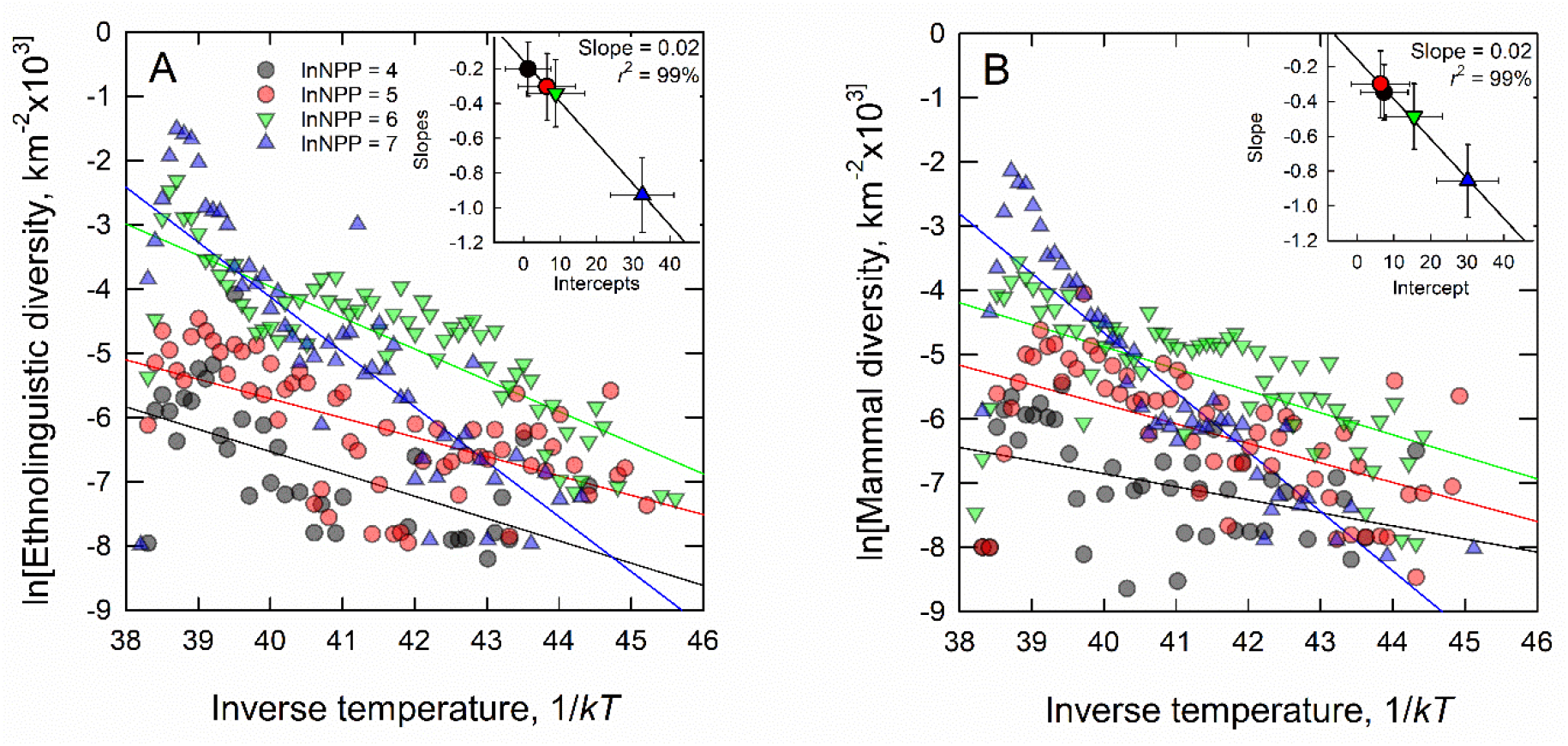
Ethnolinguistic (A) and mammal species (B) diversity by inverse temperature in bins of 1lnNPP. Inset figures are correlation plots of the slopes and intercepts per 1lnNPP bin. Solid lines are OLS regression fits. The data show the temperature dependent kinetics of diversity become systematically steeper in more productive environments.

## Discussion

Our results show that both mammal species and ethnolinguistic diversity increase superlinearly with environmental productivity at rates predicted by macroecological theory as the kinetics of diversity are faster than the kinetics of environmental productivity. This is because increasingly productive environments are also more interactive meaning that not only is more energy available, but there are increasingly different ways in which that energy is apportioned and competed over. More productive environments are also more complex, allowing more ways for diversity to be housed per unit area. To show this, we derived a simple model of diversity kinetics from well-established ecological theory from which we proposed four hypotheses whose predictions could be tested with data. Our results provide support for the four hypotheses posed in this paper, suggesting both mammal and ethnolinguistic diversity are positively density-dependent.

Our first hypothesis proposed that diversity is a function of kinetics. Indeed, for both mammal species and ethnolinguistic populations diversity increases with temperature exponentially at rates consistent with ecological kinetics (Figure 3B) [54]. As such, a linear increase in temperature leads to a multiplicative increase in diversity and the rate of this increase is well-predicted by ecological theory indicating that the rate of increase in diversity is consistent with the overall kinetics of ecosystem metabolism.

Our second hypothesis predicted that kinetics of diversity is faster than the kinetics of environmental productivity. This is because the kinetics of diversity are predicted to be both a function of the kinetics of productivity, and the kinetics of interaction rates. Results show that this prediction is supported by data where the kinetics of both mammal and ethnolinguistic diversity - *E*_*M*_ = 0.70 and *E*_*E*_ = 0.71, respectively - are both significantly faster than the kinetics of productivity, *E*_*P*_ = 0.52 .

Our third hypothesis predicted the scaling of diversity and productivity to be superlinear. Data show that for both mammal species and cultures diversity scales superlinearly with environmental productivity (for mammals *β*_*M*_ = 1.23 and for cultures *β*_*L*_ = 1.40), and the scaling exponent is consistent with the predicted ratio of kinetic terms (Figure 3B). This superlinearity describes increasing returns to scale in diversity with environmental productivity; for mammalian species a doubling of environmental productivity leads to ~23% more diversity than would be predicted by environmental energy availability alone, and in human cultures this increase is ~40%. The exact mechanisms that drive these increases are the subject of ongoing work.

Our fourth hypothesis predicted that as environmental productivity increases the temperature-dependence of the diversity response should also increase (Figures 4A and 4B). This result may be the strongest demonstration of the kinetic hypothesis as it shows even when holding environmental productivity constant, warmer environments are not only more diverse than less productive environments, but the rate of the increase in diversity is progressively faster in more productive environments. Richer environments house more species that interact with each other at increasingly faster rates. Therefore, mammalian and cultural diversity are both consistent with Red Queen dynamics driven by the temperature-dependence of interaction rates [72].

This biogeographic convergence of biological and cultural diversity is striking given the differences in the evolutionary mechanisms and time scales of biological and cultural evolution [73,74]. Perhaps most obviously, by definition, mammal species diversity is interspecific whereas ethnolinguistic diversity is intraspecific. Mammalian evolution has a time depth of many millions of years while human languages evolve on the scale of hundreds to thousands of years. Moreover, the pathways and currencies of biological and cultural evolution differ [75–77], and so an *a priori* explanation of why these different evolutionary mechanisms should converge on similar biogeographic distributions is not obvious from the dynamics of the evolutionary mechanisms themselves.

While biological and cultural evolutionary processes might be similar in principle [34,65,78,79] and even share similar mathematical representations, the detailed mechanisms of the two evolutionary processes are entirely different. Biological evolution is a statistical consequence of the differential transmission of genetic and epigenetic information over generations in response to environmental selective constraints that impact the ability of individuals to compete for finite resources and produce viable offspring. The evolutionary timescale of a biological species is commonly on the order of a few million years [80]. Cultural evolution is a statistical consequence to the differential transmission of information conveyed between individuals through the medium of language at many scales of social organization via multiple transmission pathways in both space and time [65,66,79]. Languages, cultures, and economies turnover at much faster time scales the biological species, from the near instantaneous to thousands of years [73,81]. One implication of the results shown here is that response of diversity to ecological kinetics is independent of specific evolutionary mechanism (i.e., genes vs. culture) otherwise cultural diversity would likely have evolved very differently in space to biological diversity. It may be the case that large-scale geological, climatic, and environmental constraints impact speciation and extinction processes in both systems in similar ways across gradients of environmental productivity, despite the different evolutionary inheritance mechanisms and time scales.

Whatever the specific similarities or differences between biological and cultural evolution the general mechanism of convergence proposed here is kinetic. In general, more productive environments are warmer than less productive environments, and with increasing temperature not only is there more energy available but there are increasing rates of interaction, including competition, specialization, and mutualisms, all of which combine to increase diversity above levels predicted by environmental productivity. Diversity increases superlinearly with environmental productivity because more productive environments are also more complex.

## Author contributions

All authors contribute equally to this work.

## Materials and methods

### Temperature and environmental productivity data

We used the WorldClim database for mean annual temperature in °C and mean annual precipitation in mm yr^−1^ at 10 arc min resolution (version 1.4; http://www.worldclim.org/bioclim). WorldClim provides interpolated global climate data for stations with 10 to 30 years of data [68]. We used estimates of annual net primary productivity (g C m^−2^ yr^−1^) from an average of 17 global models at a spatial resolution of 0.5 degrees [69]. All data were first projected with the WGS 1984 Cylindrical Equal Area projection and then processed across our language polygons using the Zonal Statistics tool in ArcGIS. The Global Self-consistent, Hierarchical, High-resolution Shoreline dataset was used as the basemap [70].

### Mammal and ethnolinguistic data

Ethnolinguistic data were downloaded from the Ethnologue, and centroids (in decimal degrees of latitude and longitude) were recorded for each individual language polygon (*N* = 7,627). Terrestrial mammal data were downloaded from PanTHERIA and centroids (in decimal degrees of latitude and longitude) were recorded for each species (*N*=5,089). Average temperature, annual precipitation, and net primary productivity were the extracted for each ethnolinguistic population and mammal species. Data layers were compiled in ArcGIS and then analyzed statistically in R [71]. We compiled a GIS of data layers for each of the variables used in the paper.

### Sampling procedure and statistics

Measuring diversity requires a sampling strategy. We use a statistical sampling technique designed to isolate the independent variable and reduce spatial autocorrelation. First, we divided the surface of the planet into a grid of 0.5 degrees of latitude by 0.5 degrees of longitude, extracted the non-oceanic cells (for a total of *n* = 56,598 cells) and calculated the available landmass per cell. For each cell we then counted the number of ethnolinguistic populations and mammal species that occur in that cell, and then divided that occurrence data by the available landmass. We also calculated the average temperature and net primary production for each cell. To sample global diversity, we then created bins of 0.1 lnNPP and 0.1 1/kT, and summed the total occurrence of ethnolinguistic and mammalian species densities, thus providing measures of the diversity per bin of independent variable controlling for the available landmass within that bin. This sampling approach has several important statistical advantages. First, we aggregate the data into bins by small increments of the independent variable, and then take statistics over the bin. This technique has the advantage of using the degrees of freedom available from the data set to estimate statistical moments within bins, and therefore isolates the effect of the independent variable on the dependent variable. Statistical error is thus minimized when estimating scaling exponents. Second, neither mammal species nor ethnolinguistic groups are statistically independent as they both share evolutionary ancestries, usually captured by phylogenetic models and a spatial structure resulting in autocorrelation. The binning procedure minimizes both forms of autocorrelation as data points are aggregated from non-local populations collected from around the planet and averaged; the residuals of the binned data are no longer correlated. Thus data used in the scaling analyses are aggregates and are statistically independent. Third, the binning method minimizes the potential weighting biases of heteroscedasticity and unequal sampling across the data set.

## Supplementary Information

### Detailed statistical Results

Statistical results below are presented by the sequence (figures) they occur in the paper.

**Figure 3A.**
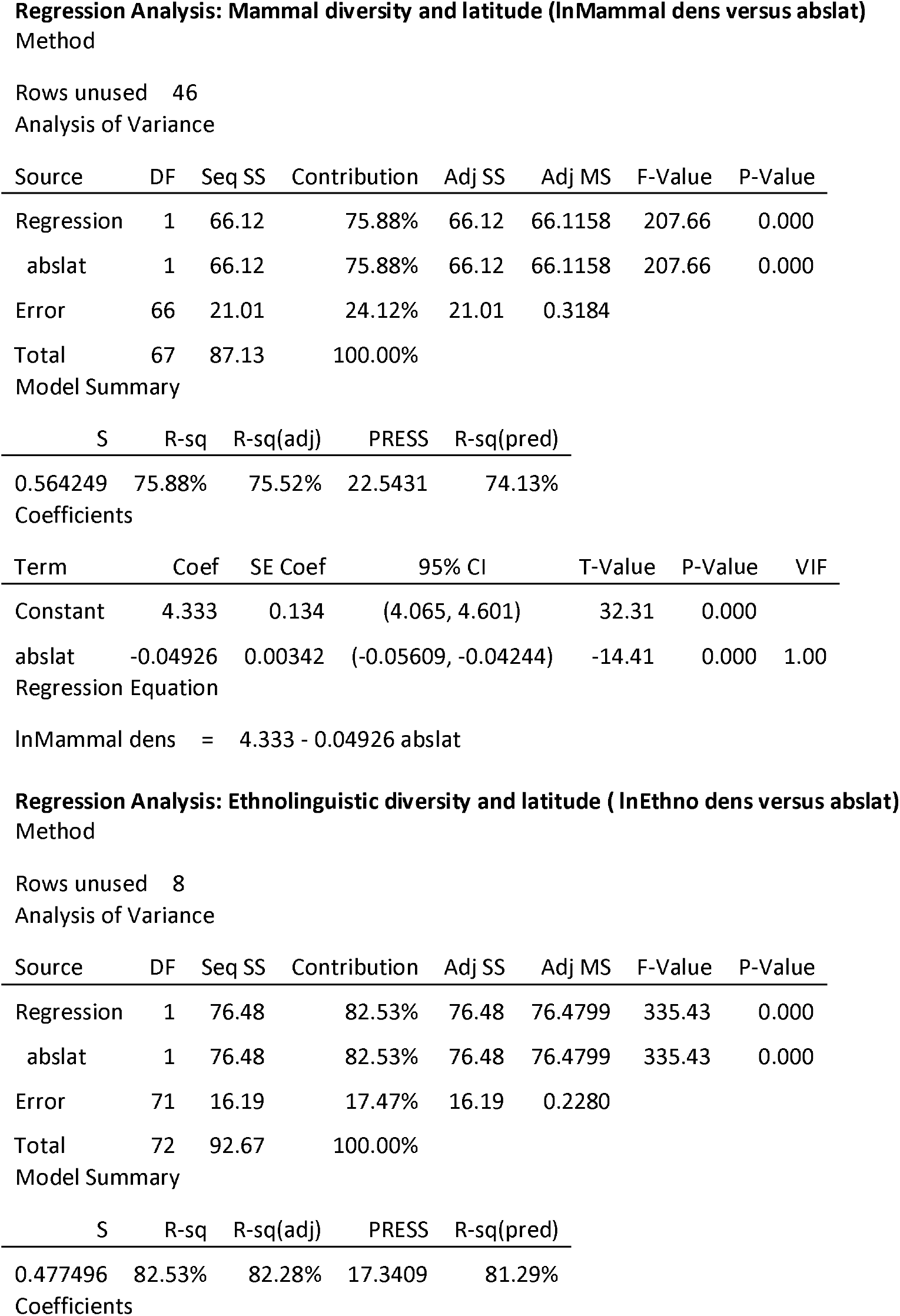
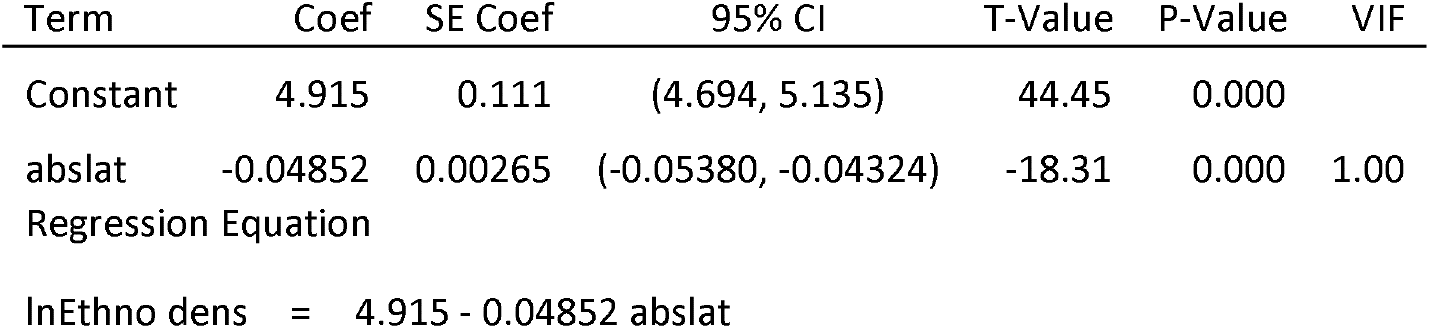

**Figure 3B.**
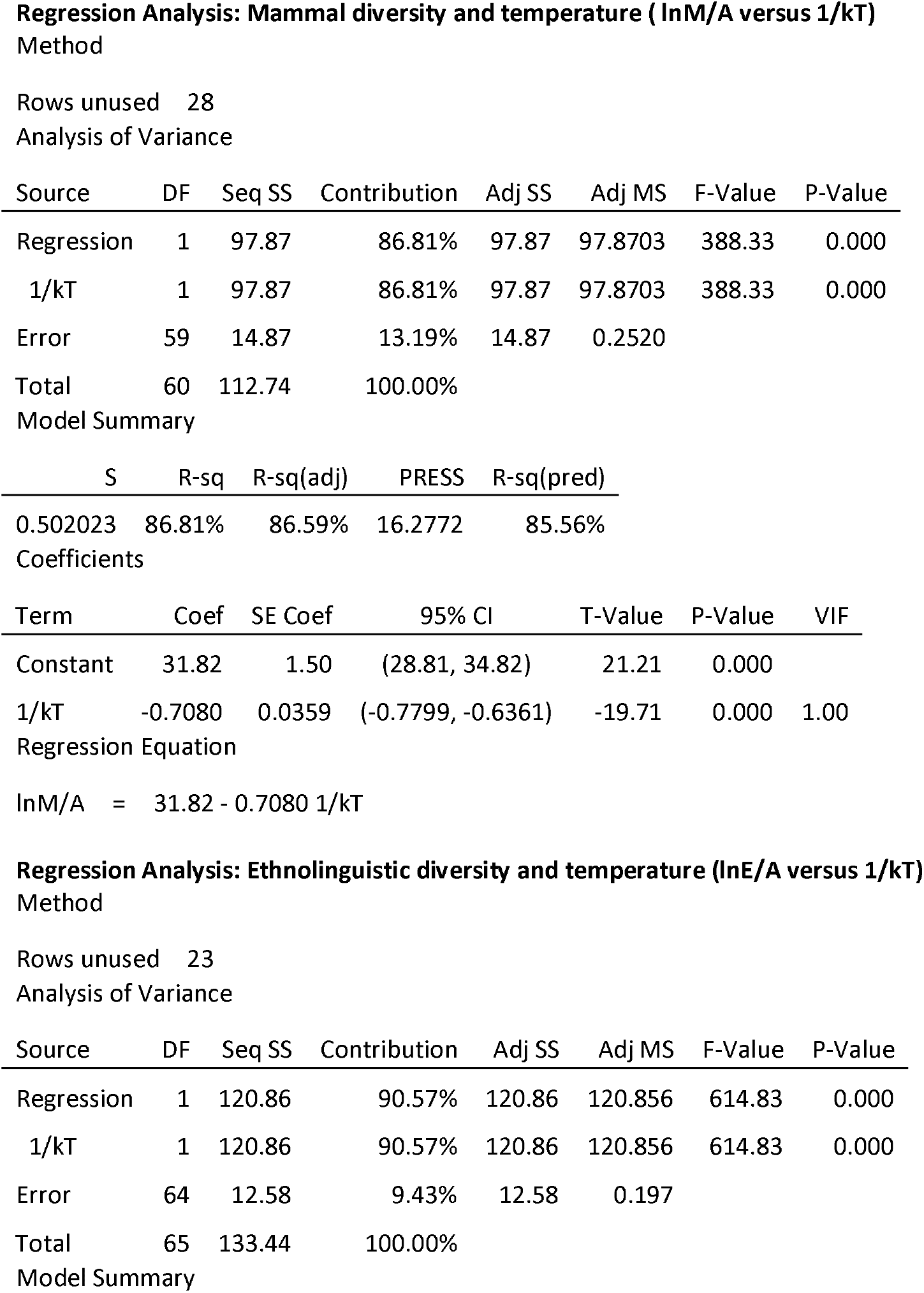
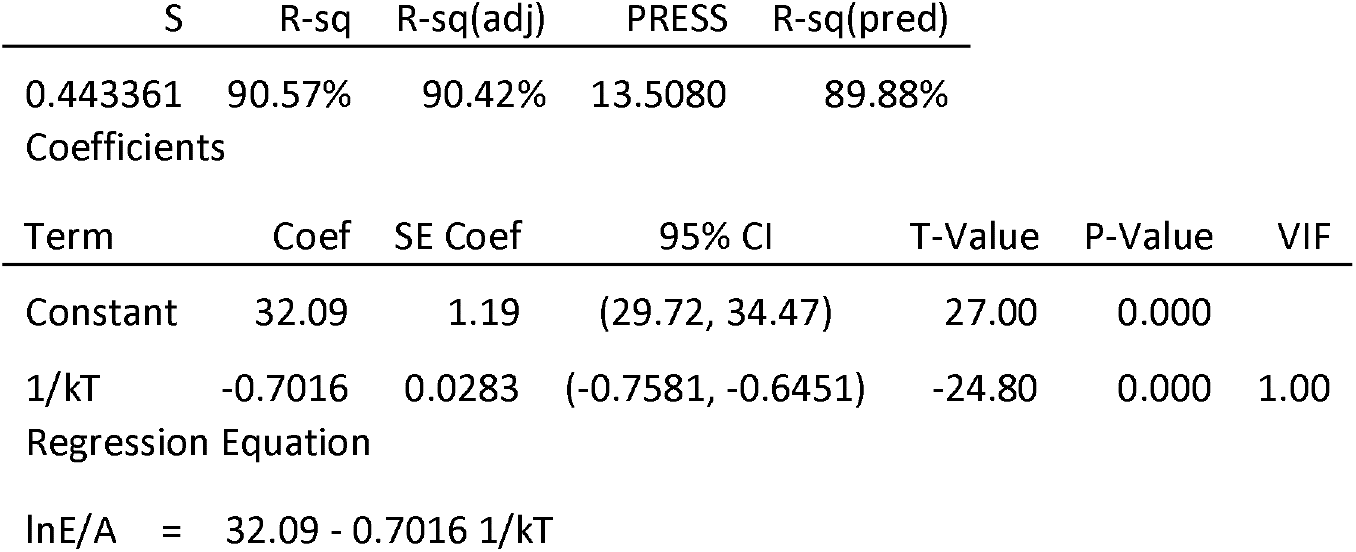

**Figure 4B.**
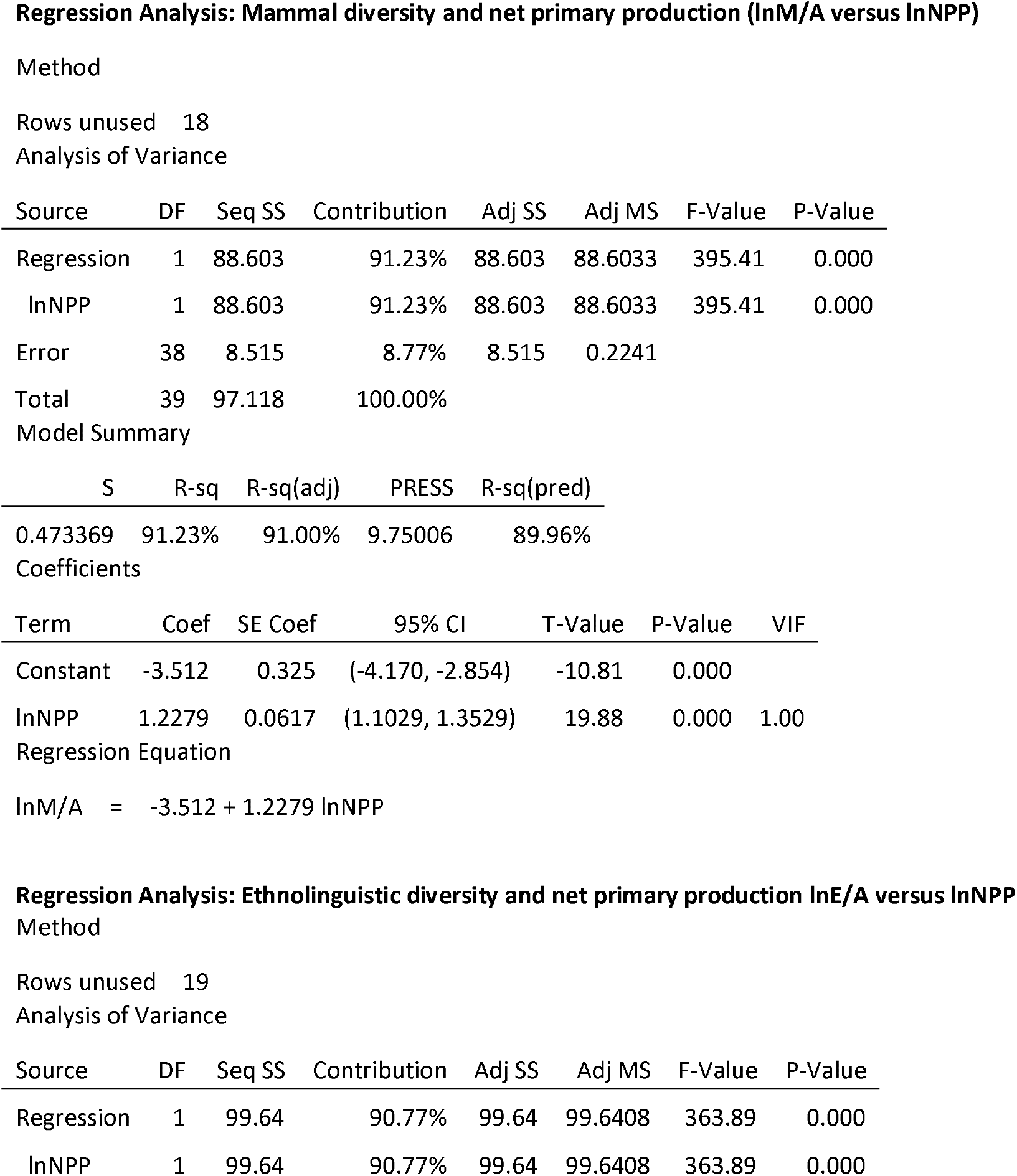
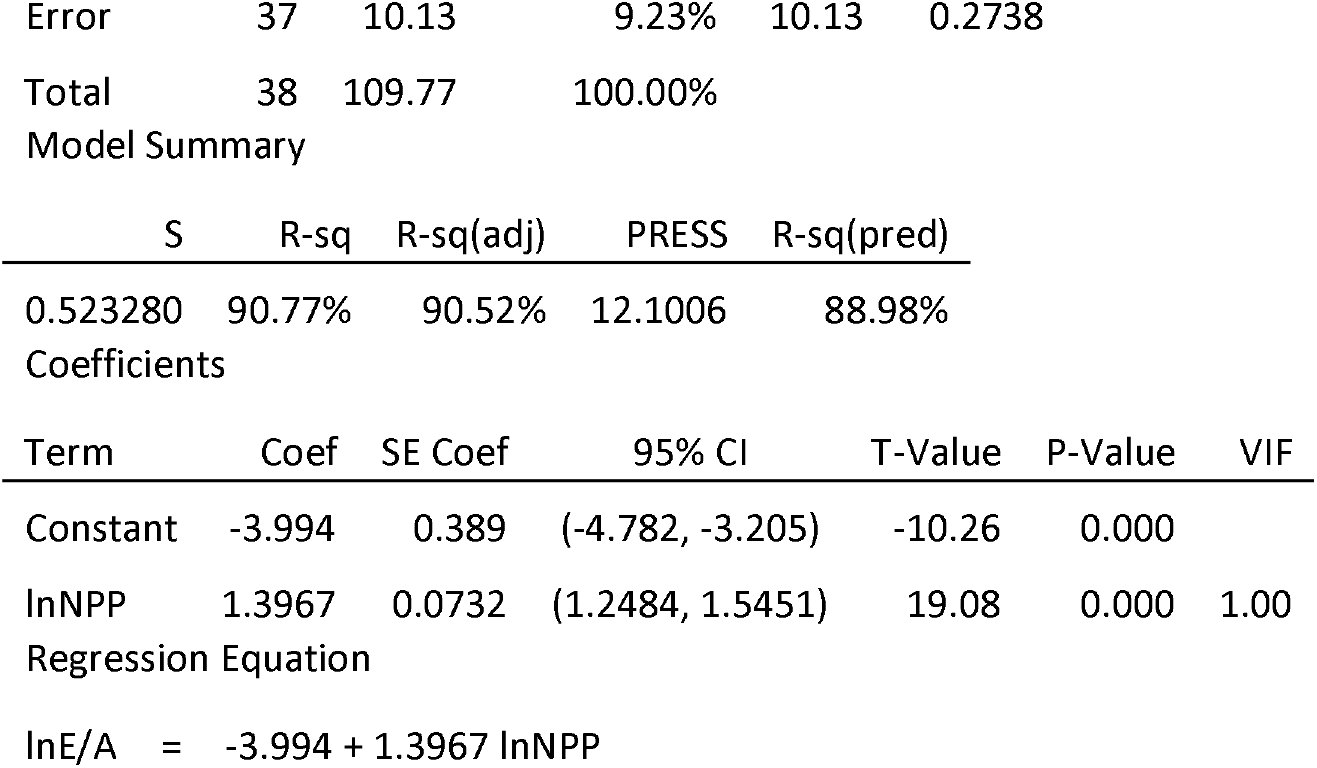

**Figure 5A.**
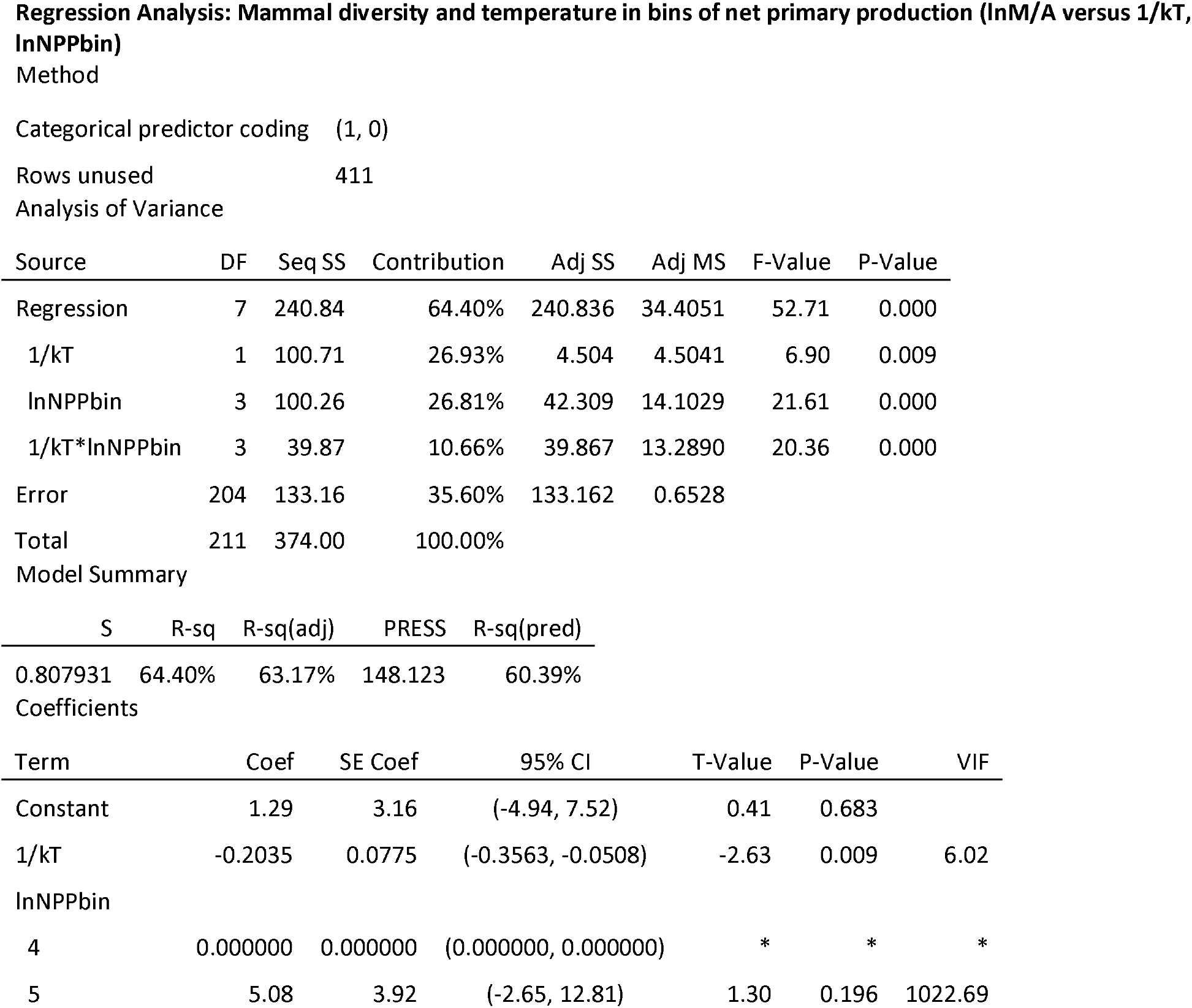
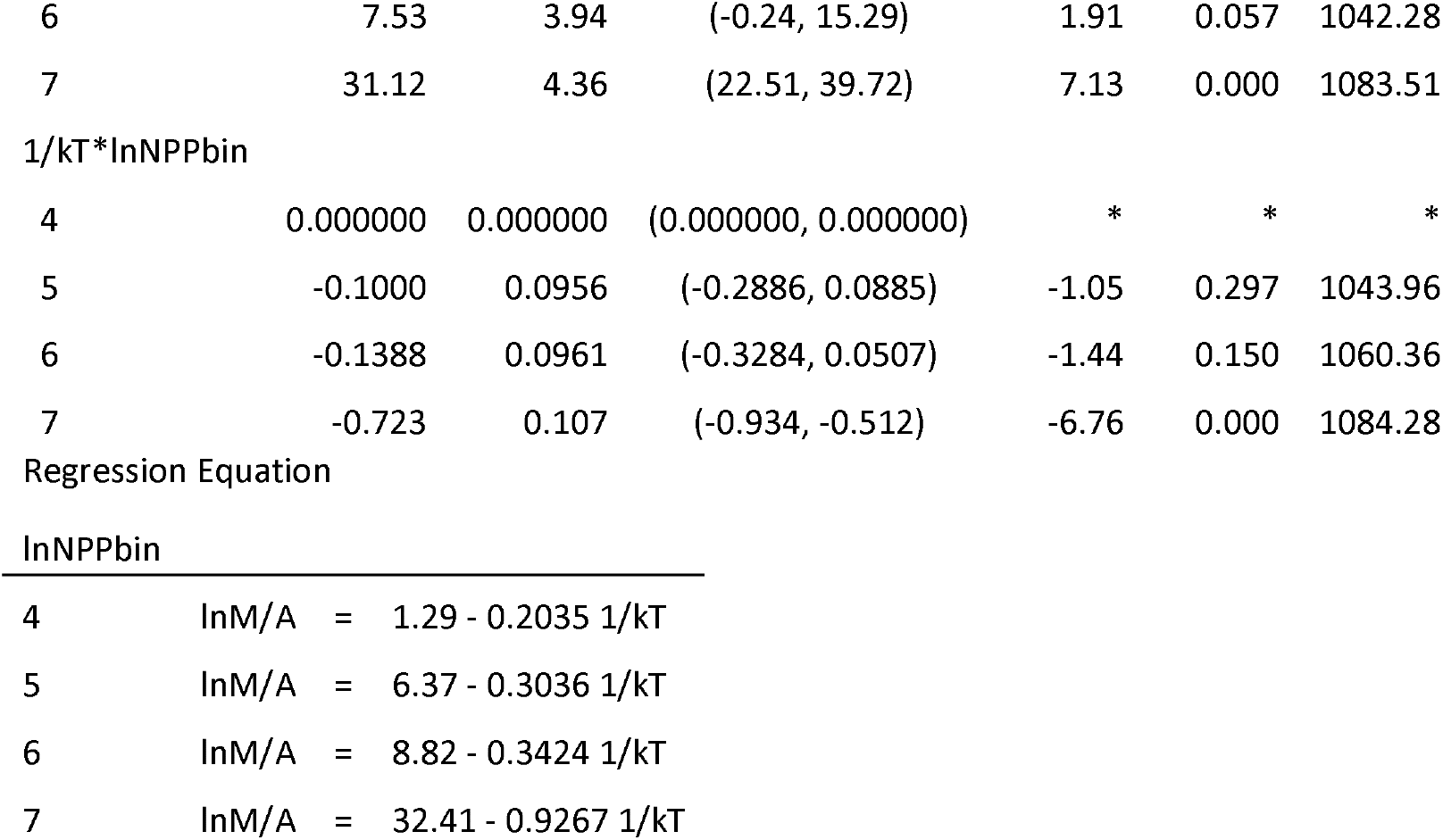

**Figure 5B.**
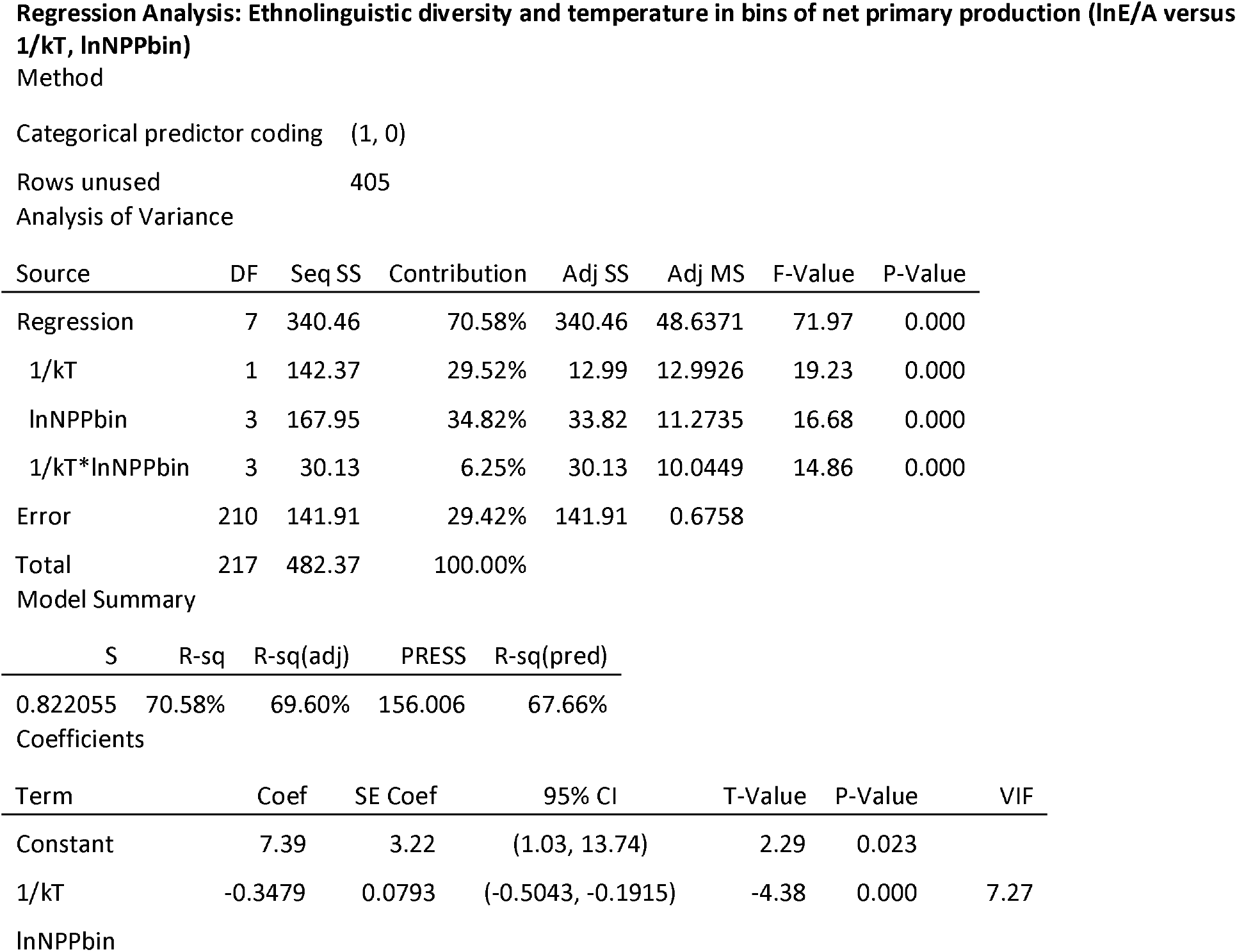
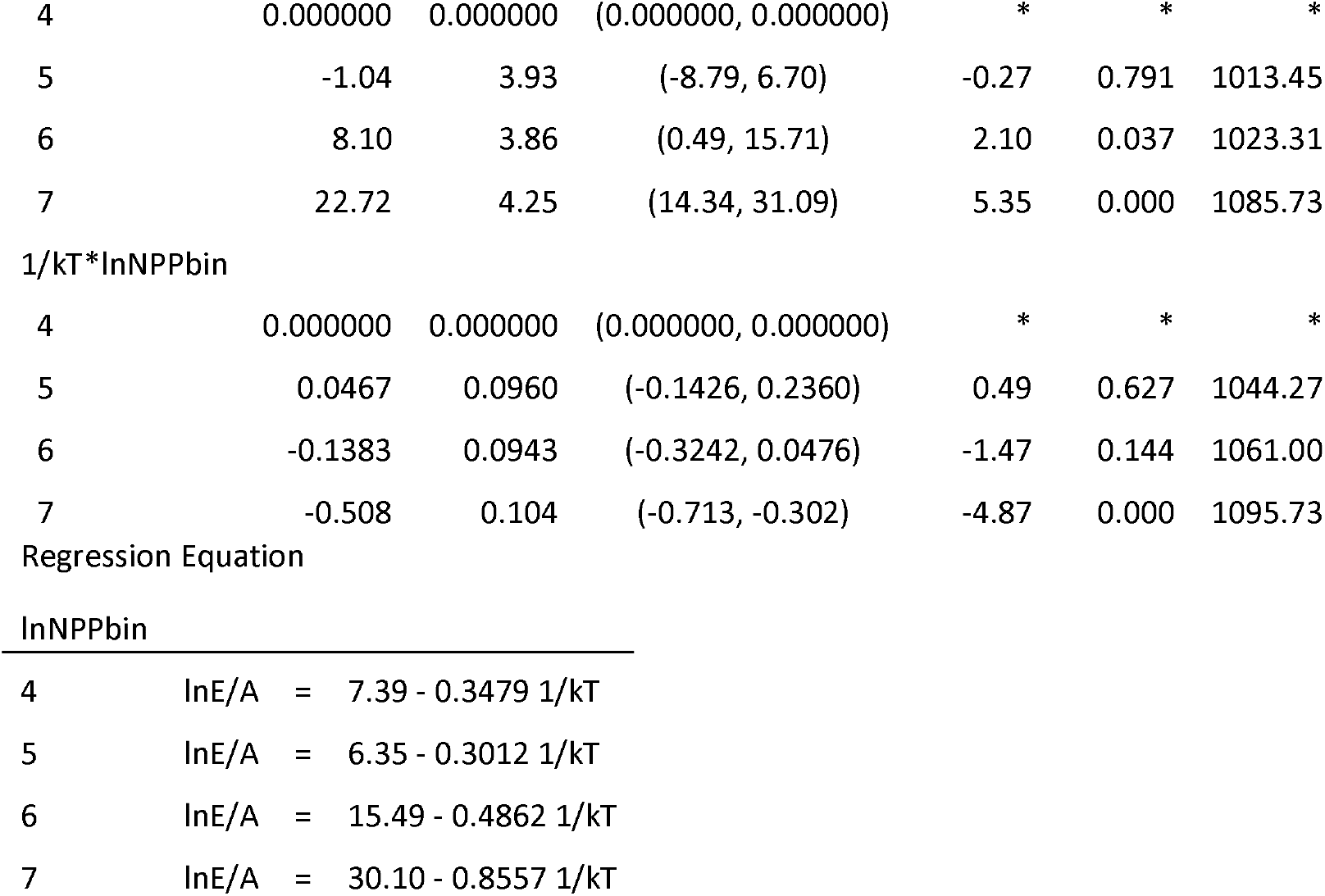

## Notes

### Competing Interest Statement

The authors have declared no competing interest.

